# CRISPR/Cas9 gene editing in the heterotrophic protist *Acanthamoeba castellanii*

**DOI:** 10.64898/2026.07.28.741101

**Authors:** Dudley Chung, Sari Matar, John M. Archibald

## Abstract

Members of the amoebozoan genus *Acanthamoeba* are unicellular heterotrophic protists that inhabit a wide range of terrestrial and aquatic environments. They are opportunistic pathogens known to host disease-causing bacteria as well large DNA viruses. While *Acanthamoeba* is commonly used to study cellular locomotion, phagocytosis, and disease, genetic tools for precise editing of the *Acanthamoeba* genome are still in their infancy. Here we have developed CRISPR/Cas9 tools for genetically engineering the model acanthamoebid *Acanthamoeba castellanii* strain Neff. In proof-of-principle experiments, we targeted the myosin-II heavy chain gene. Evidence for editing was found using reverse transcriptase PCR, microscopy, and DNA sequencing, which also identified unique alleles for the target locus. Collectively, we demonstrate that CRISPR can be employed to edit *Acanthamoeba* genes using a knock-in approach. This work serves as a foundation to further develop *A. castellanii* as a model system with which to study diverse questions in cell and molecular biology, biochemistry and evolution.

## Introduction

*Acanthamoeba* species are free-living unicellular, heterotrophic eukaryotes found in soil and diverse aquatic habitats, both natural and artificial.^1-7^ They feed on other microorganisms such as bacteria, some of which use *Acanthamoeba* as a host or vector to protect against environmental stresses. A prominent example is the bacterium *Legionella pneumophila*, which causes Legionnaires’ disease in humans. Research on *Acanthamoeba castellanii* has shown that *L. pneumophila* can replicate within a protective vacuole, and passage through which results in increased bacterial virulence.^8^

*Acanthamoeba* is itself an opportunistic human pathogen. It can cause skin lesions and sinus infections as well as *Acanthamoeba* keratitis, an eye disease that if left untreated can cause blindness.^9^ Along with other amoebae such as *Balamuthia, Acanthamoeba* spp. can also cause the fatal central nervous system disease granulomatous amoebic encephalitis.^9^ In addition to being used to study bacterial pathogenesis, *Acanthamoeba* is used as a model system to study fundamental cellular processes such as motility, differentiation, autophagy and phagocytosis.^9,10^ Despite its utility as a lab workhorse *Acanthamoeba* remains relatively understudied at the molecular biology level, in part because of its genomic complexity. The nuclear genomes of *A. castellanii* strains sequenced thus far are relatively small — approximately 45 megabase pairs in size.^11,12^ However, they exhibit complex and variable ploidy^13,14^ and have acquired bacterial and viral genes over the course of their evolution by lateral gene transfer.^11,12,15^

With the goal of expanding the molecular biology ‘toolbox’ of *A. castellanii*, we explored the feasibility of gene editing using CRISPR. In nature, the CRISPR system serves a bacterial defense system against bacteriophages; in the lab, the CRISPR-associated effector endonuclease Cas9 can be engineered to target and cut DNA in a sequence-specific manner.^16^ Here we provide a proof-of-principle demonstration of CRISPR-Cas9 for the successful editing of *A. castellanii* genes. We engineered DNA plasmids to allow us to perform CRISPR and attempted a knock-in strategy to determine gene editing efficiency. The presence of edited genes was verified using reverse transcriptase PCR (RT-PCR), microscopy, and sequencing of the target region.

## Results

### Validation of Cas9 and guide RNA expression in the delivery vectors by RT-PCR and fluorescence microscopy

We constructed a delivery plasmid containing a GAPDH promoter-driven eSpCas9 linked at the C-terminal end by monomeric green fluorescent protein (mEGFP) as well as a lysine tRNA promoter-driven gRNA scaffold (Figure 1A). The plasmid was then transformed into *A. castellanii* and gene expression of both Cas9 and gRNA was validated by RT-PCR. We observed amplicons approximately 100 base-pairs (bp) in size corresponding to the gRNA scaffold (Figure 1B). *A. castellanii* cells transformed with eSpCas9-mEGFP were deemed positive for mEGFP expression by RT-PCR and fluorescence microscopy (Figure 1C and D).

**Figure 1.**
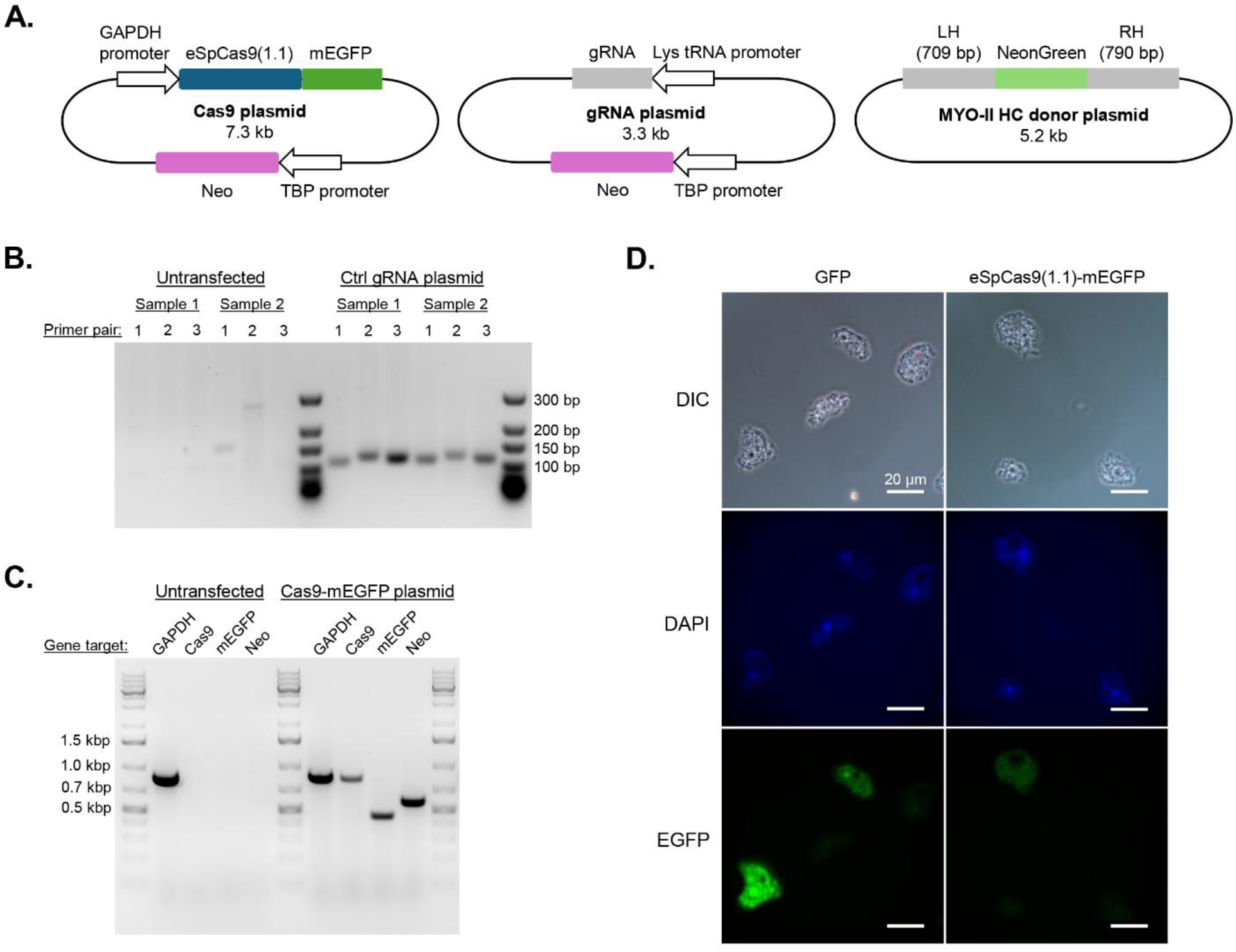
Construction and validation of a CRISPR-Cas9 system for gene editing in *Acanthamoeba castellanii*. A) Schematic diagram of the Cas9, gRNA and homology donor plasmids used in this study (LH – left homology arm, RH – right homology arm). B) RT-PCR of *A. castellanii* cells transfected with expression plasmid containing control gRNA. Extracted RNA was reverse transcribed and PCR amplified using three different primer sets that bind to the crRNA (gRNA + scaffold RNA). C) RT-PCR of *A. castellanii* cells transfected with expression plasmid containing Cas9-mEGFP. Extracted RNA from two different transfected samples were reverse transcribed and PCR amplified using primer sets that bind to the Cas9 and mEGFP coding sequences. RT-PCR of Neomycin and GAPDH regions served as controls. D) Representative fluorescence microscopy images showing GFP fluorescence in *A. castellanii* cells transfected with pGAPDH-EGFP and the Cas9-mEGFP expression plasmid.

### CRISPR editing of Myosin-II Heavy Chain

Myosin-II is a motor complex involved in actin remodelling. It consists of two identical heavy chains and two pairs of light chains, and can self-assemble to form bipolar filaments.^17-19^ In *Acanthamoeba*, myosin-II has been shown to be distributed in the cytoplasm, but appears concentrated in the cell cortex as well as at the cleavage furrow during cytokinesis.^19^ We used CRISPR-Cas9 to attach the monomeric green fluorescent protein NeonGreen to the N-terminal end of myosin-II heavy chain protein (NG-MYO-II HC).^20^ Our experiments showed NG-MYO-II HC localization patterns consistent with other studies using antibody labeling and transient transfection.^19,21^ More specifically, while fluorescent signal was clearly observed in the cytoplasm, we also saw enhanced fluorescence at the trailing end of motile *A. castellanii* cells, in agreement with previous reports suggesting that myosin-II provides constriction forces (Figure 2A and B).^22^

**Figure 2.**
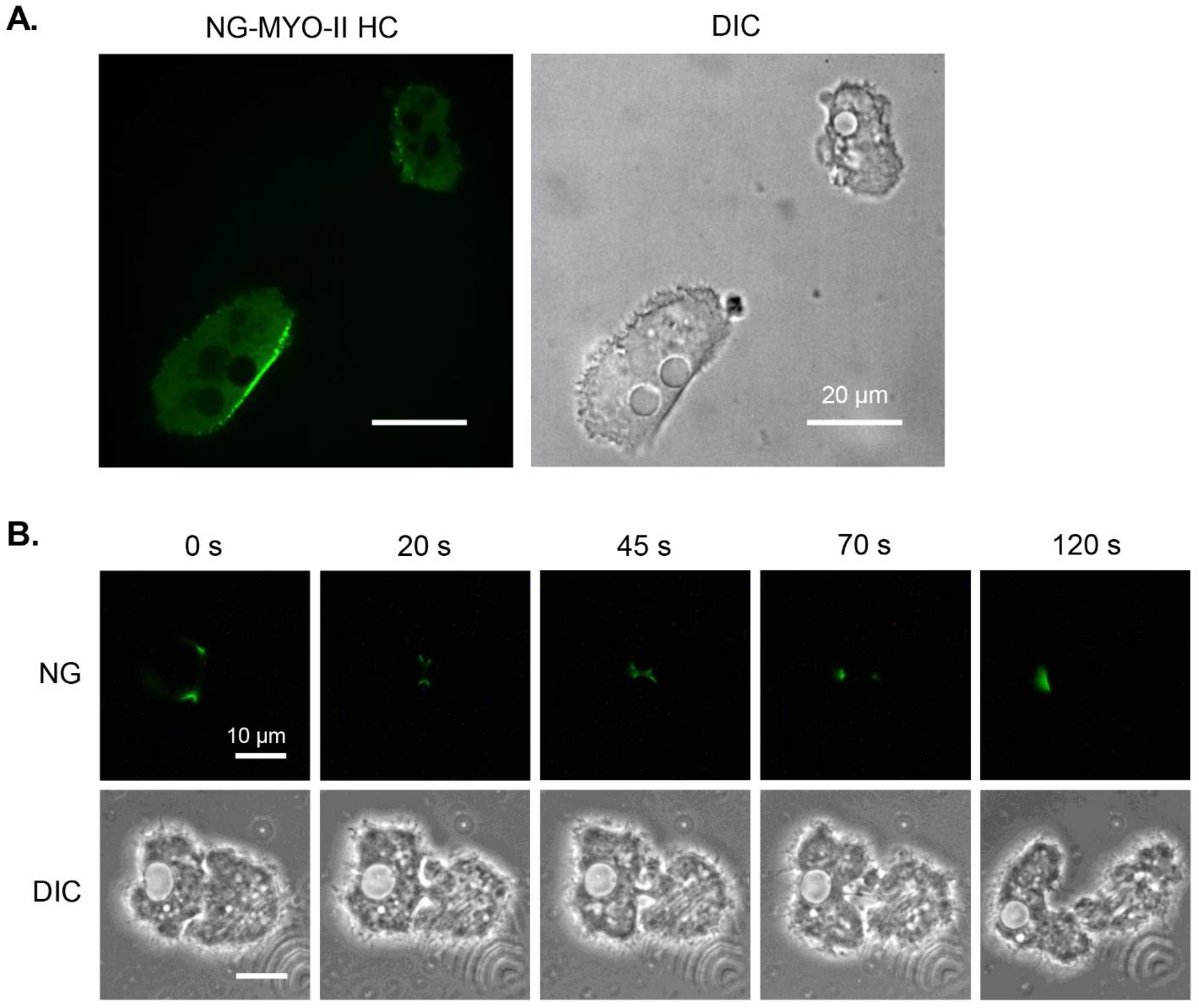
CRISPR knock-in of NeonGreen into the myosin-II heavy chain gene of *Acanthamoeba castellanii* shows expected protein localization patterns. A) Representative image of CRISPR-edited cells showing localization of NeonGreen-myosin-II heavy chain (NG-MYO-II HC) to the trailing edge of locomotive cells. B) Representative timelapse images of a dividing cell showing that NG-MYO-II HC concentrates at the cleavage furrow during cytokinesis.

Sanger sequencing of PCR amplicons of the N-terminal coding region of MYO-II HC in *A. castellanii* strain Neff confirmed CRISPR gene editing (Figure 3). However, a mixture of edited and unedited alleles was found in the recovered population of cells, indicating incomplete editing. In addition, we found allelic diversity, with at least two variants of the MYO-II HC gene present in the population. These results are supported by the close matching of PCR- and Sanger-derived sequences obtained herein with Illumina sequences generated for whole genome sequencing.^12^ Sections of the region sequenced herein map to one or the other existing published scaffolds, i.e., Neff-v1 and Neff-v2 (Figure 3B and C). This suggests that the MYO-II HC gene sequence variants in the published *A. castellanii* strain Neff genome assembly are an amalgamation of multiple distinct alleles.

**Figure 3.**
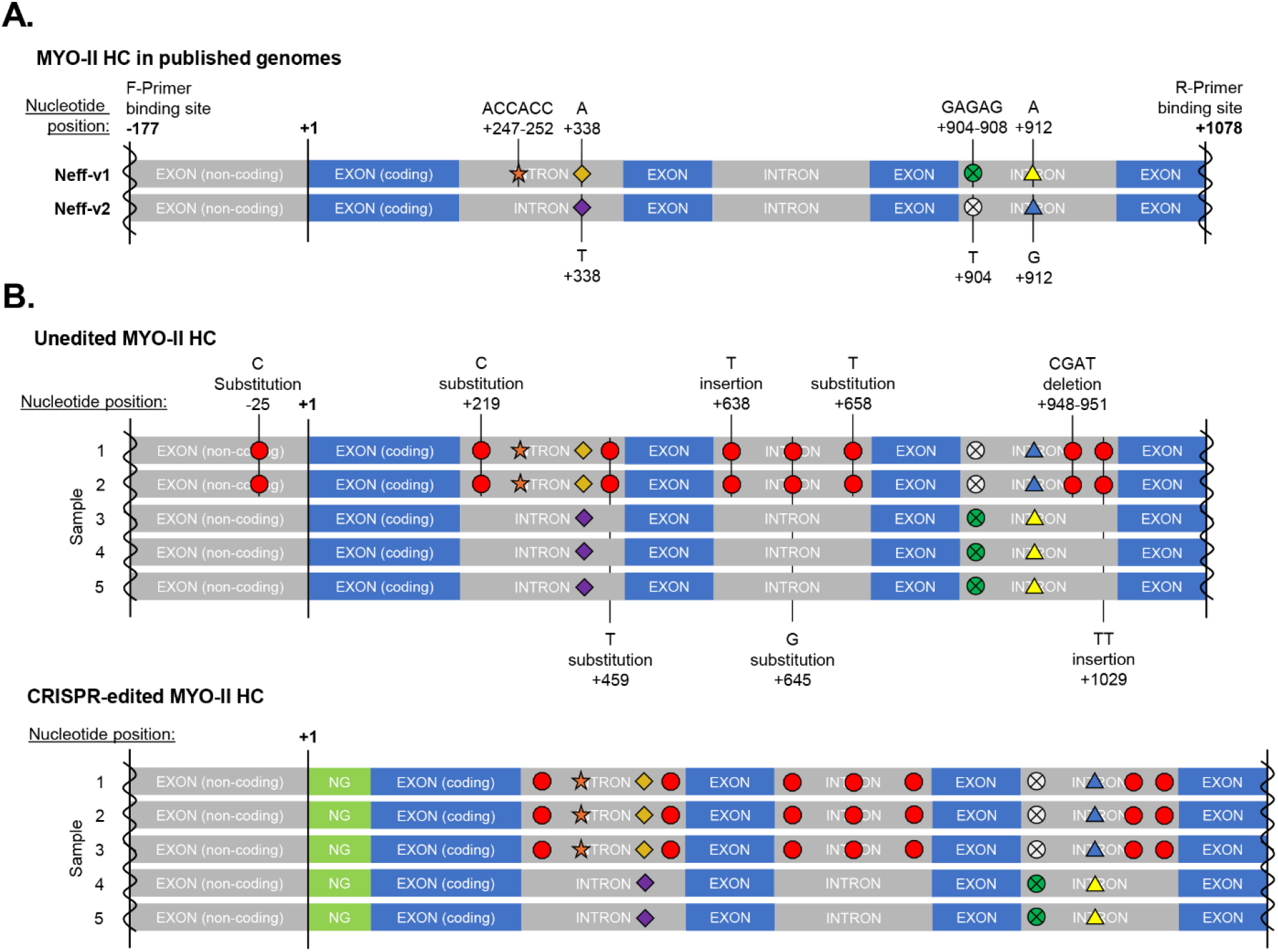
Sequencing of MYO-II HC gene PCR products from *Acanthamoeba castellanii* confirms CRISPR editing and reveals unique allelic patterns. A) Schematic illustration showing nucleotide-level differences between the Neff-v1 and Neff-v2 scaffolds for the N-terminal coding region of the myosin-II heavy chain gene. The position of the 3’ nucleotide on both the forward and reverse sequencing primer is indicated. B) Schematic illustration of the nucleotide-level differences at the same region of individual clonal isolates of un-transfected *Acanthamoeba* and ones transfected with the CRISPR construct.

## Discussion

*Acanthamoeba castellanii* Neff and related strains have long been used to study cellular motility, host-pathogen interactions, and cellular processes such as phagocytosis.^9,10^ Using gene-editing technologies such as CRISPR can greatly complement such laboratory studies. We have demonstrated that CRISPR-mediated insertion of DNA sequences into *A. castellanii* genes is possible. We used this technique to incorporate NeonGreen to the N-terminus of Myosin-II HC and have generated a stable cell line.

*Acanthamoeba* contains additional types of myosin motor proteins, each with presumably different roles.^22^ Concurrent temporospatial investigation of the other types of myosin motor proteins inside the same cell is possible using CRISPR. While our results show positive gene editing in *A. castellanii*, complete and uniform allelic editing was not achieved. This is presumably due to the polyploid nature of the genome, which is still being investigated.^12,13^ At the present time, the CRISPR tools generated herein are most useful for knock-in experiments and the tracking of cellular and molecular phenotypes via endogenous tagging.

During the execution of our study, uniform editing of the *A. castellanii* genome was demonstrated by Bisio *et al*., who integrated a Cas9-gRNA unit into their cells by incorporating homology arms on each end of the Cas9-gRNA.^23,24^ The incorporation of the Cas9-gRNA sequence by homology-directed repair to generate the knock-out mutant allows for continuous expression of Cas9-gRNA to edit unmodified alleles. The authors also demonstrated CRISPR editing of the genome of a giant virus that infects the amoeba, a powerful tool for dissecting host-virus interactions. In the system developed herein, the expression vectors lack homology arms resulting in a lower likelihood of integration. It is also conceivable that the plasmid may no longer be retained once selection pressure (i.e., antibiotic selection) is removed; therefore, overall expression levels of the Cas9 protein is likely more transient. This may be beneficial in situations where gene dosage matters and altering every copy of a gene causes unintended phenotypic effects, such as disease development in multicellular organisms due to haploinsufficiency or triplo-sensitivity. Continued development of molecular tools for genome editing in *A. castellanii* and other amoebozoans will make it possible to interrogate diverse aspects of the biology of these complex and important microorganisms.

## Methods

### Plasmid Construction

The tRNA-gRNA scaffold, Cas9-EGFP, Neomycin resistance (Neo), and homology donor sequence were cloned into the pGEM-T vector (Promega #A3600) and propagated in JM109 competent *E. coli* (Promega #L2005). Positive selection of transformants was achieved by plating on LB agar with 100 µg/mL ampicillin. *A. castellanii* tRNAs were predicted using tRNAscan-SE and ARAGORN.^25,26^ The Cas9 from eSpCas9(1.1) (Addgene plasmid #71814) was codon optimized at the N- and C-terminal ends for *A. castellanii* using synthesized gene fragments (Integrated DNA Technologies) and joined using overlap PCR.^27^ Guide RNA sequences (underlined) were designed using the gRNA design software in Geneious; Control gRNA F – GGCT<u>ACAATCCGCTGCGGCTGACG</u>; Control gRNA R – AAAC<u>CGTCAGCCGCAGCGGATTGT</u>; MYO-II HC gRNA F – GGCTA<u>AGATGGCCGCCCAGCGCAGG</u>; MYO-II HC gRNA R – AAAC<u>CCTGCGCTGGGCGGCCATCT</u>T. The gRNA scaffold sequence was modified as described in Chen *et al*., 2013 and contains two *Bbs*I restriction sites for inserting the gRNA sequence.^28^ The *Acanthamoeba* GAPDH promoter and the TBP-Neomycin cassette were PCR amplified from pGAPDH-EGFP as described in Bateman, 2010.^29^ The homology donor sequence, consisting of a codon-optimized NeonGreen and 700-800 nucleotides flanking the starting codon on myosin-II HC, were synthesized as gene fragments (Integrated DNA Technologies) and combined using overlap PCR.

### Cell Culture

*Acanthamoeba castellanii* strain Neff (ATCC 30010) was cultured at room temperature (20-22 °C) in PYG medium (0.75% yeast extract, 0.75% peptone, 2 mM potassium phosphate monobasic (KH_2_PO_4_), 1 mM magnesium sulfate heptahydrate (MgSO_4_•7H_2_O)) containing 1.5% (w/v) D-glucose, and supplemented with 0.1mM ferric citrate, 0.05 mM CaCl_2_, 1 μg/mL thiamine, 0.2 μg/mL D-biotin, and 1 ng/mL cobalamin.

### Transfection

Cells were transfected as described previously using Superfect (Qiagen #301305) or calcium phosphate.^30,31^ For calcium phosphate transfection, a 150 µL mixture of 250 mM CaCl_2_ and a 5:3:1 ratio (by µg of plasmid) of donor template, gRNA, and Cas9 plasmid (e.g., 13.33 µg of donor template, 8 µg of gRNA vector, 2.67 µg of Cas9 vector), was mixed dropwise with 2x HeBS buffer (274 mM NaCl, 10 mM KCl, 1.4 mM Na_2_HPO_4_, 15 mM D-glucose, 42 mM HEPES, pH 7.10). The final mixture was thoroughly distributed into a 6-well plate containing 2.75×10^5^ adherent cells grown in 1 mL PYG medium and allowed to incubate for 4 h at room temperature with gentle rocking. Cells were exposed to 10% glycerol prepared in HEPES buffer for 3 min, then removed and replaced with PYG medium. Cells were treated with 5 µg/µL G418 (InvivoGen #ant-gn-1) 24 h after transfection, which was increased to 25 µg/µL once transfectants have recovered. Enriched populations of edited *A. castellanii* cells were obtained by serial dilution and cell picking.

### RNA Extraction and RT-PCR

Total RNA was obtained using TRIzol™ extraction following the manufacturer’s protocol (Thermo Fisher #15596026). cDNA was generated using the Maxima H Minus First Strand Synthesis kit (Thermo Fisher #K1681) using site-specific primers. The forward primer used amplify the Control guide RNA from cDNA is F – ACAATCCGCTGCGGCTGACG, which is paired with either reverse primer R1 – CGACTCGGTGCCACTTTTTCAAGTTG, R2 – AAAAAAGCACCGACTCGGTGCCAC, or R3 – GCACCGACTCGGTGCCACTTTTTC. PCR primers used to amplify GAPDH are F – TGTCGCACATCAAGGTCGGTAT, and R – GAGGAGTGGATGAAGTCGGAGG. Primers to amplify Cas9 are F – ATCGTGTGGGATAAGGGCCG, and R – CCGGTGATGCTCTGGTGGAT. PCR primers to amplify EGFP are F – TTCTTCAAGTCCGCCATGCC, and R – TCGTCCATGCCGAGAGTGAT. Primers used to amplify Neo are F – AAGATGGATTGCACGCAGGTTC, and R – TCCACCATGATATTCGGCAAGC.

### Genomic DNA Extraction

*A. castellanii* cells were lysed using Stan’s Extraction Buffer (200 mM Tris-HCl pH 7.5, 250 mM NaCl, 25 mM EGTA, 0.5% SDS) and treated with 0.05 mg/mL RNase A (37 °C for 30 min) then with 1 mg/mL Proteinase K (55 °C for 1 h). Total genomic DNA was isolated by performing two rounds of phenol:chloroform:isoamyl alcohol (25:24:1) (Sigma-Aldrich #77617) extraction, then two rounds of chloroform:isoamyl alcohol (24:1) extraction before precipitation using isopropanol. DNA was washed twice using 80% ethanol and resuspended in 10 mM Tris-HCl, pH 8.0. Confirmation that the gene locus was edited was achieved by Sanger sequencing (GENEWIZ) of agarose gel-purified PCR amplicons (Qiagen #28704); Forward myosin-II HC primer – CCCCAAGCACCAAGAAGAGGA, Reverse myosin-II HC primer – CTCCGTCTTACCAGCTCCCG.

### Microscopy

Cells grown on No. 1.5 coverslips were fixed for 15 min at room temperature with 4% formaldehyde prepared in Page’s amoeba saline (2.1 mM NaCl, 16 µM MgSO_4_**·**7H_2_O, 41 µM CaCl_2_**·**2H_2_O, 1.0 mM Na_2_HPO_4_, 1.0 mM KH_2_PO_4_). Samples were washed with PAS, DAPI stained (300 mM) and mounted with ProLong™ Diamond antifade mountant (Thermo Fisher #P36965). Fixed cells were imaged with an Olympus BX43 inverted microscope equipped with an X-Cite Series 120Q fluorescence lamp (EXFO Photonic Solutions Inc.). Locomotive cells were imaged on a Zeiss Marianas confocal spinning disk microscope equipped with a 488 nm Argon laser.

## Acknowledgements

This work was supported by a Gordon and Betty Moore Foundation grant (GBMF5782) awarded to JMA. We thank Feng Zhang for providing the eSpCas9(1.1) plasmid. We also thank Graham Dellaire (Dalhousie University) for providing training and technical assistance on using the confocal spinning disk microscope.

